# Genome wide identification of bacterial genes required for plant infection by Tn-seq

**DOI:** 10.1101/126896

**Authors:** Kevin Royet, Nicolas Parisot, Agnès Rodrigue, Erwan Gueguen, Guy Condemine

## Abstract

Soft rot enterobacteria (*Dickeya* and *Pectobacterium*) are major pathogens that cause diseases on plants of agricultural importance such as potato and ornamentals. Long term studies to identify virulence factors of these bacteria focused mostly on plant cell wall degrading enzymes secreted by the type II secretion system and the regulation of their expression. To identify new virulence factors we performed a Tn-seq genome-wide screen of a transposon mutant library during chicory infection followed by high-throughput sequencing. This allowed the detection of mutants with reduced but also increased fitness in the plant. Virulence factors identified differed from those previously known since diffusible ones (secreted enzymes, siderophores or metabolites) were not detected by this screen. In addition to genes encoding proteins of unknown function that could be new virulence factors, others could be assigned to known biological functions. The central role of the FlhDC regulatory cascade in the control of virulence was highlighted with the identification of new members of this pathway. Scarcity of the plant in certain amino acids and nucleic acids required presence of the corresponding biosynthetic genes in the bacteria. Their products could be targets for the development of antibacterial compounds. Among the genes required for full development in chicory we also identified six genes involved in the glycosylation of the flagellin FliC, glycosylation, which in other plant pathogenic bacteria contributes to virulence.

**Author summary:** Identification of virulence factors of plant pathogenic bacteria has relied on the test of individual mutants on plants, a time-consuming method. New methods like transcriptomic or proteomic can now be used but they only allow the identification of genes induced during the infection process and non-induced genes may be missed. Tn-seq is a very powerful method to identify genes required for bacterial growth in their host. We used for the first time this method in a plant pathogenic bacteria to identify genes required for the multiplication of *Dickeya dadantii* in chicory. We identified about 100 genes with decreased or increased fitness in the plant. Most of them had no previously described role in bacterial virulence. We unveiled important metabolic genes and regulators of motility and virulence. We showed that *D. dadantii* flagellin is glycosylated and that this modification confers fitness to the bacteria during plant infection. Our work opens the way to the use of Tn-seq with bacterial phytopathogens. Assay by this method of large collections of environmental pathogenic strains now available will allow an easy and rapid identification of new virulence factors.

## Introduction

*Dickeya* are broad-host range phytopathogenic bacteria belonging to the Pectobacteriaceae family [1] that provoke the soft rot disease on many plant species. They are the cause of important losses on economically important crops such as potato, chicory and ornamentals. Identification and studies on the virulence factors of these bacteria have been performed mostly on the model strain *D. dadantii* 3937 and focused mainly on three domains/aspects, known to be important for disease development: plant cell wall degrading enzymes, the type III secretion system and iron metabolism [2]. Secretion of plant cell wall degrading enzymes has long ago been identified as the bacteria main virulence factor. Many studies focused on the identification and characterization of these secreted enzymes, mostly pectinases [3], of the regulators controlling their production (*kdgR, pecS, pecT, hns, gacA*), [4–8] of the genes whose expression is coregulated with that of the secreted enzyme genes [9, 10], and of the mechanism of their secretion by the type II secretion system [11]. Although of a lesser importance for *Dickeya* virulence, the same type of approach has been used to identify type III secretion system regulators and effectors [12] [13] [14]. Moreover, struggling for iron within the plant is strong. *D. dadantii* acquires this metal through production of two siderophores, chrysobactin and achromobactin [15] [16] [17]. Omics approaches have also been used to identify genes induced during plant infection [18] [19] [20]. These studies now provide a clearer picture on a complex network of factors required for *D. dadantii* virulence [2, 21]. However, these approaches may have missed some important factors not targeted by these analyses. More global screens need to be performed to identify these factors. Libraries of transposon-induced mutants were tested on plants to find mutants showing reduced virulence with *Pectobacterium carotovorum* and *atrosepticum*, two other soft rot enterobacteria [22–24]. These studies identified auxotrophs, mutants defective in production or secretion of exoenzymes and in motility. Other mutants with a more complex phenotype were not characterized at this time. Moreover, the number of tested mutants was limited by the necessity to test individually each mutant on plant. This type of work has never been performed on *Dickeya* strains. To have a more complete view of the genes required for the virulence of *Dickeya*, we used a high-throughput sequencing of a saturated transposon library (Tn-seq) to screen tens of thousands random insertion mutants of *D. dadantii* in laboratory medium and during infection of chicory. Tn-Seq involves creating large transposon libraries, growing the mutants in a control and a selective condition, sequencing the transposon insertion sites with next-generation sequencing, mapping sequence reads to a reference genome and comparing the number of read in each gene in the two conditions. Tn-seq has been extensively used to uncover essential genes required for mouse colonization by human pathogens *Vibrio cholerae* [25], *Pseudomonas aeruginosa* [26] and *Streptococcus pneumoniae* [27] or plant root colonization by *Pseudomonas simiae* [28] or multiplication of *Pantoea stewartii* in corn xylem [29]. This latter bacteria relies on the massive production of exopolysaccharides to block water transport and cause wilting. Thus, Tn-seq is a very powerful method to identify genes required for bacterial growth in their host. By applying this technique to screen a *D. dadantii* mutant library in chicory, we identified metabolic pathways and bacterial genes required by a necrotrophic bacteria for growth *in planta*. Among them, we found a cluster of genes required for flagellin glycosylation, a modification known to be important for several plant pathogenic bacteria virulence.

## Results and discussion

### Characterization of *D. dadantii* 3937 *Himar1* transposon library

Many tools are available to perform Tn-seq [30]. In order to perform a Tn-seq experiment with *D. dadantii* 3937, we used a *Himar9* mariner transposon derivative carrying MmeI restriction sites in the inverted repeats (IR) and a kanamycin resistance cassette between the IRs [31]. We carried out a biparental mating between *E. coli* and *D. dadantii* on M63 agar medium without carbon source and amino acids. We obtained approximately 300 000 colonies that were pooled. Subsequent DNA sequencing (see below) showed the presence of transposon insertions in amino acid, vitamin, purine or pyrimidine biosynthesis pathways, demonstrating that mating on M63 minimal medium does not prevent the obtention of auxotroph mutants. To identify essential genes, mutants were grown in LB medium for several generations. Two DNA libraries were prepared from two cultures and subjected to high-throughput sequencing. The mariner transposon inserts into TA dinucleotides. The TPP software [32] was used to determine the number of reads at each TA site for each biological replicate. *D. dadantii* genome has 171,791 TA sites that can be targeted by the *Himar9* transposase. Pairs of biological replicates were compared. 37,794 and 48,101 unique insertions in TAs were detected in each sample, which corresponds to 22 and 28% density of insertion respectively (Table 1). The average number of reads per TA is 88 and 75, respectively. The results were reproducible with a Pearson correlation coefficient of 72% (Fig. S1) The location of the unique insertions showed an even distribution around the chromosome (Fig. 1A). For each gene, we calculated a log_2_ fold change (FC) corresponding to a ratio between the measured number of reads and the expected number of reads. The density plot (Fig. 1B) indicates that essential and non-essential genes are easily distinguishable, confirming the good quality of our Tn-seq libraries.

**Fig. 1.**
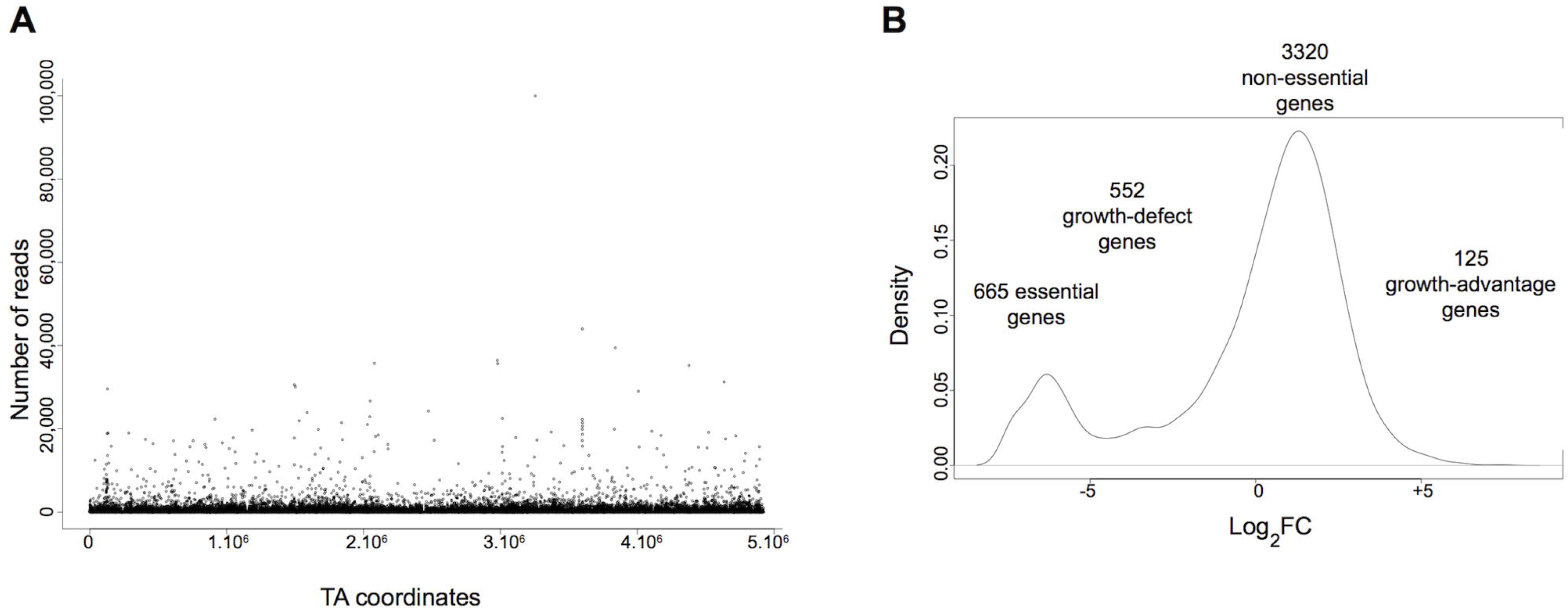
Quality control of the Tn-seq *D. dadantii* 3937 libraries. (A) Frequency and distribution of transposon sequence reads across the entire *D. dadantii 3937* genome. The localization of transposon insertions shows no bias throughout the genome of *D. dadantii 3937*. B) Density plot of log_2_FC (measured reads/expected reads per gene).

**TABLE 1.** Tn-Seq analysis of *Dickeya dadantii* 3937

| Mutant pool | Total no. of reads | No. of reads containing Tn end | No. of reads normalized<sup>a</sup> | No. of mapped reads to unique TA sites | No. of mapped reads to unique TA sites after LOESS correction | Density (%)<sup>b</sup> | Mean read count per TA<sup>c</sup> |
| --- | --- | --- | --- | --- | --- | --- | --- |
| LB #1 | 23,152,186 | 22,647,343 | 18,748,028 | 13,166,770 (70 %) | 12,904,900 (69 %) | 28 % | 75 |
| LB #2 | 30,105,412 | 27,963,154 | 18,748,028 | 15,535,291 (83 %) | 15,195,582 (81 %) | 22 % | 88 |
| Chicory #1 | 18,925,029 | 18,748,028 | 18,748,028 | 17,535,146 (94 %) | 14,906,888 (79 %) | 24 % | 87 |
| Chicory #2 | 27,607,717 | 26,555,297 | 18,748,028 | 17,477,706 (93 %) | 16,955,724 (90 %) | 23 % | 99 |

<sup>a</sup> The number of reads containing the sequence of a Tn end were normalized for each sample according to the Chicory #1
<sup>b</sup> *Dickeya dadantii* 3937 genome has 171,791 TA sites. The density is the % of TAs for which mapped reads has been assigned by the TPP software.
<sup>c</sup> The mean value of mapped reads per TA

Then, gene essentiality of the Tn-seq input libraries was determined by using the TRANSIT software [32]. We decided to use the Hidden Markov Model (HMM) which predicts essentiality and non-essentiality for individual insertion sites since it has been shown to give good prediction in datasets with density as low as 20% [32]. The HMM analysis led to the identification of 665 genes essential for growth in LB (ES), representing 14% of the genes of *D. dadantii* 3937, a number in the range of those found for this type of analysis with bacteria. The transposon we used does not allow us to discriminate between the direct effect of the insertion or a polar effect on downstream genes. Goodall et al [33] have shown that this overestimates the number of essential genes. Thus 665 must be considered has an upper limit of the number of essential genes.

552 genes were categorized as Growth Defect genes (GD, i.e. mutations in these genes lead to loss of fitness), 125 as growth advantage genes (GA, i.e mutations in these genes lead to gain of fitness) and 3320 as non-essential genes (NE) (Table S5 and Fig. 1B).

#### Genes necessary for chicory leaf maceration

We used chicory leaf infection as a model to identify *D. dadantii* genes required for growth in plant tissues. Biological duplicates were performed to insure the reproducibility of the results. Each chicory was inoculated with 10^7^ bacteria from the mutant pool and after 2 days more than 10^10^ bacteria were collected from the rotten tissue. Sequencing transposon insertion sites in these bacteria followed by the TPP analysis indicated a density of unique insertion in TAs comparable to that of the input datasets (23–24%). Surprisingly, the results were more highly reproducible than in LB with a very high Pearson correlation coefficient of 98% (Fig. S1).

In order to test the statistical significance of the identified genes conferring to *D. dadantii* a loss or a gain of fitness *in planta*, we performed the RESAMPLING (permutation test) analysis of the TRANSIT software. The RESAMPLING method is a variation of the classical permutation test in statistics that sums the reads at all TA sites for each gene in each condition. It then calculates the difference of the sum of read-counts between the input (LB) and output (chicory) datasets. The advantage of this statistical method is to attribute for each gene an adjusted p-value (q-value). Genes with a significant difference between total read-counts in LB and chicory achieve a q-value ≤ 0.05. The method also calculates a log_2_ fold-change (log_2_FC) for each gene based on the ratio of the sum of read counts in the output datasets (chicory) versus the sum of read counts in the input (LB) datasets [32]. Applied to our Tn-seq datasets and selecting only genes achieving a FDR adjusted p-value (q-value) ≤ 0.05, we identified 122 genes out of 4666 required for fitness *in planta*, as shown with the volcano plot of RESAMPLING results comparing replicates grown in LB versus *in planta* (Fig. S2). For these 122 genes, we applied an additional cutoff by removing 20 genes with a mean read count in LB <5 (less than 5 reads in average / TA). These genes were categorized as ES or GD in LB. We also removed from the analysis 6 genes with a log_2_FC comprised between −2 and 2. By applying all these criteria, we retained only 96 genes for a further analysis (Table 2). 92 of them were identified as GD genes in the chicory (log_2_FC ≤2), the 4 left as GA genes in the chicory (log_2_FC ≥2). A possible polar effect for genes being part of an operon is analysed in Table 2: if a GD gene is upstream of another GD gene in the same operon, a polar effect of insertions in the first gene on the second one cannot be excluded. Some of these genes, in bold in Table 2, were already known to play a role in *D. dadantii* virulence, confirming the validity of the Tn-seq approach. Using the Kyoto Encyclopedia of Genes and Genomes (KEGG) [34], we discovered that certain metabolic pathways and biological functions are very important for growth in chicory (Table S4). We highlight some of them in the next sections of the article.

**TABLE 2.**
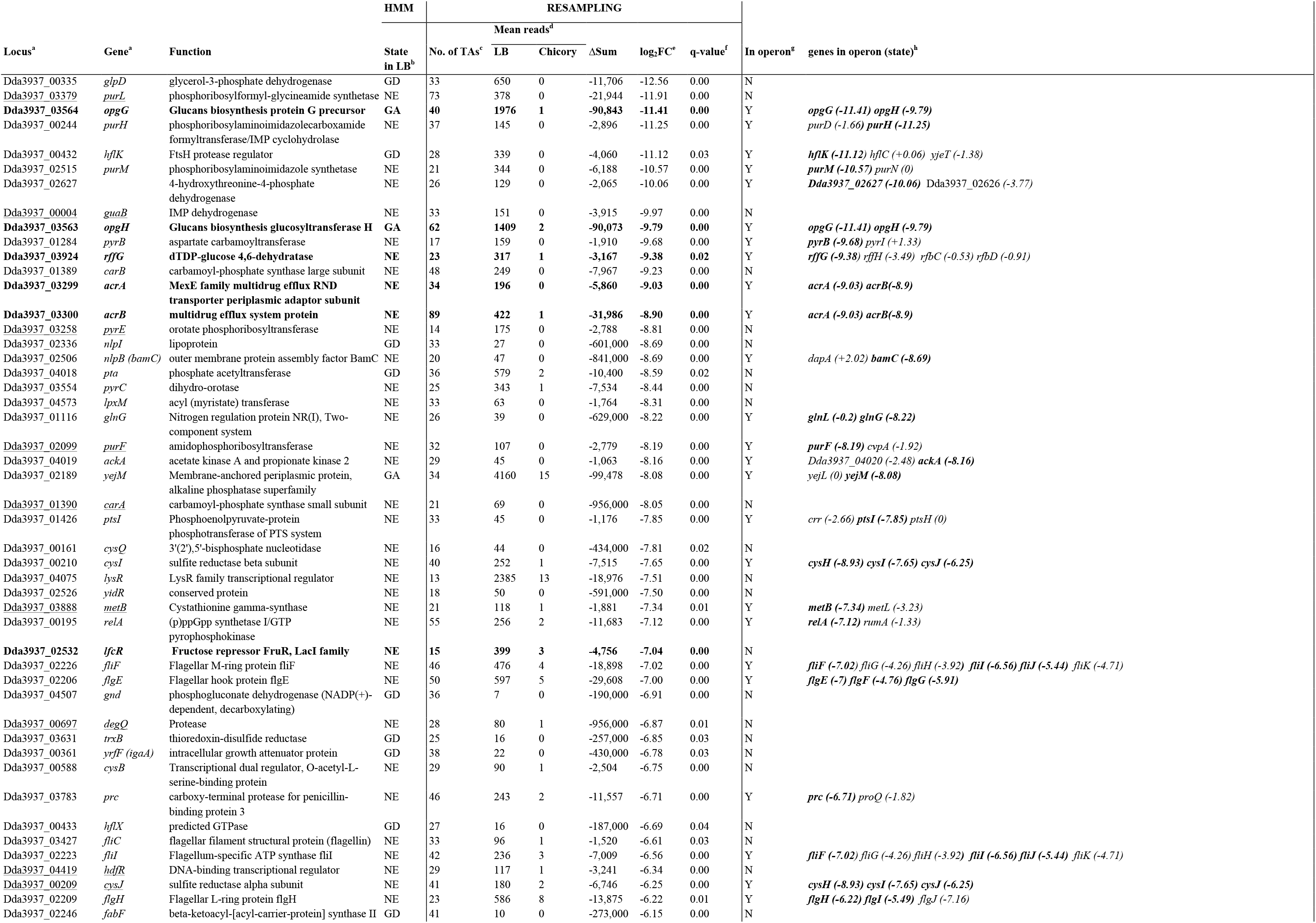
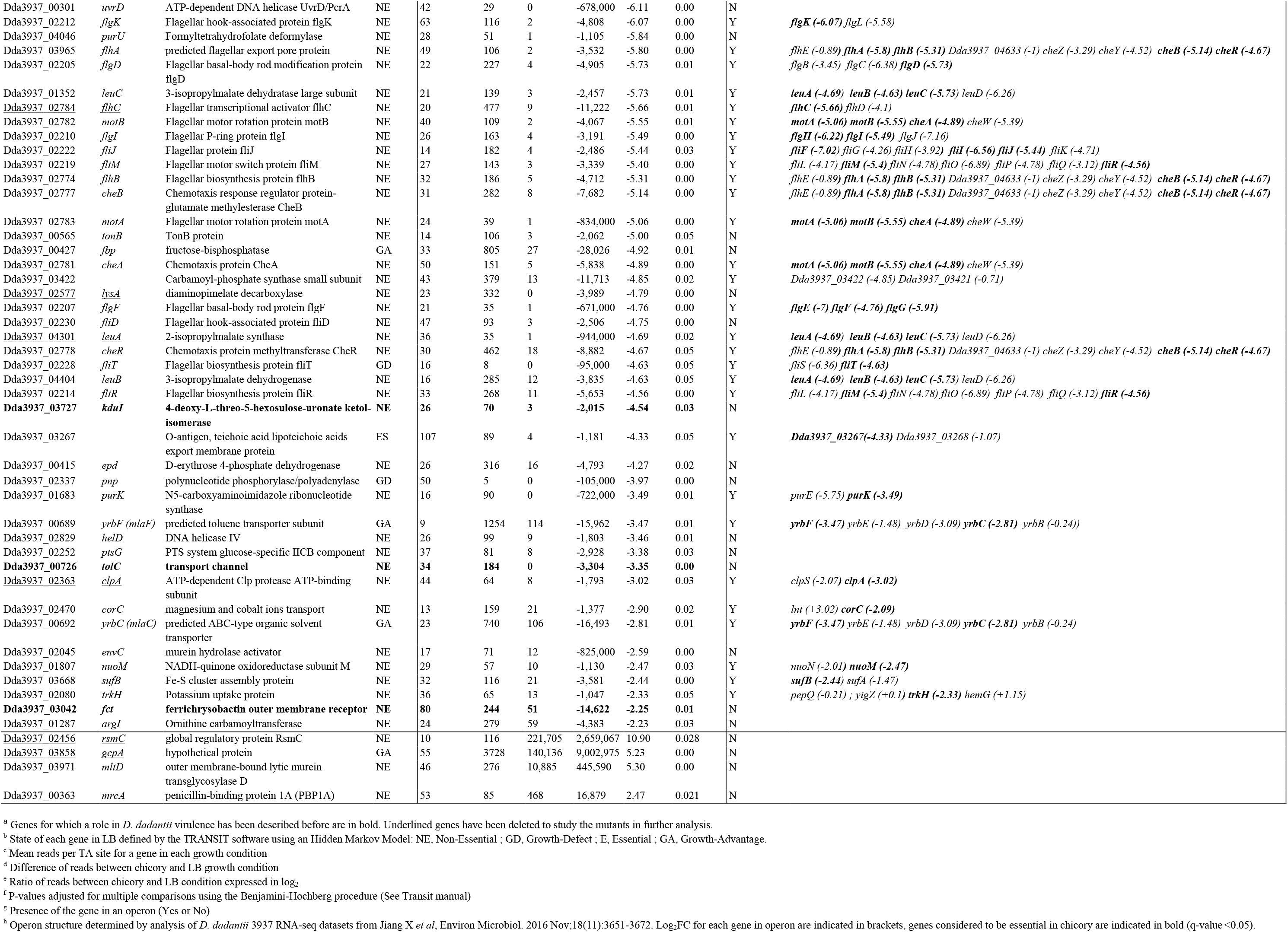
Genes identified by Tn-seq exhibiting a fitness change in chicory compared to LB.

### Analysis of the genes of *D. dadantii* required for plant colonization

#### (i) Metabolism

Chicory appears as an environment in which amino acids, nucleic acids and some vitamins (pyridoxal) are scarce. Of the 92 genes identified as GD genes *in planta*, 8 are involved in purine and 7 in pyrimidine metabolisms (Table S4). In the purine metabolism pathway, the inosine monophosphate (IMP) biosynthesis pathway that produces IMP from L-glutamine and 5-phosphoribosyl diphosphate is particularly important for *D. dadantii in planta* since 5 out of the 10 genes of this pathway are significantly GD genes *in planta* (Fig. 2). IMP is the precursor of adenine and guanine. Next, IMP can be converted in xanthosine 5′-phosphate (XMP) by the IMP dehydrogenase GuaB. *guaB* gene is also a GD gene *in planta*, with a strong log_2_FC of −10.06 (Fig. 2). In the pyrimidine synthesis, the uridine monophosphate (UMP) biosynthesis pathway that converts L-glutamine to UMP, a precursor of uracyl, is very important *in planta* since *carAB, pyrB, pyrC* and *pyrE*, involved in this enzymatic pathway, are all required for growth *in planta* (Fig. 2). This pyrimidine biosynthesis pathway is specific to bacteria. It is noteworthy that in the human pathogen *S. pneumoniae*, mutants of this pathway have a fitness defect in the nasopharynx of infected mice [27]. Hence, it looks that the pyrimidine biosynthesis pathway is particularly important for multiplication of some bacterial species in the host.

**Fig 2.**
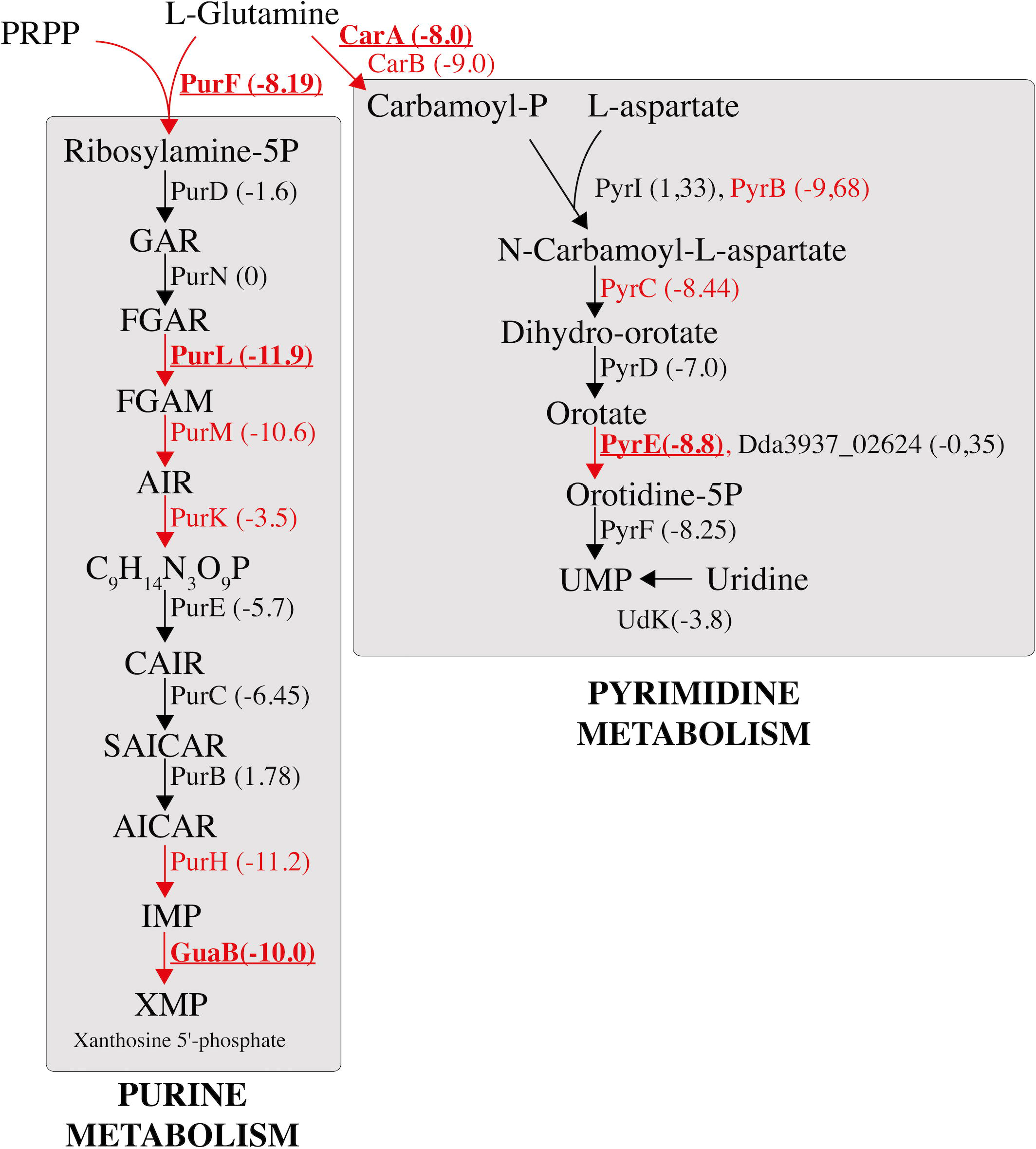
Scheme of the purine and pyrimidine biosynthesis pathways in *D. dadantii* that produce XMP (purine metabolism) and UMP (pyrimidine metabolism) from L-glutamine. In red are indicated the growth defect genes in chicory that pass the permutation test (q-value ≤ 0.05). In bold are genes for which GD phenotype was tested and confirmed with in frame deletion mutants. The log_2_FC of read numbers between chicory and LB for each gene is indicated in bracket. Some genes do not pass the permutation test (in black) but have a strong negative log_2_FC. PRPP: 5-phosphoribosyl-1-pyrophosphate; GAR: 5’-phosphoribosyl-1-glycinamide; FGAM: 5’-phosphoribosyl-*N*-formylglycinamide; AIR: 5′-phosphoribosyl-5-aminoimidazole; CAIR: 5’-phosphoribosyl-5-aminoimidazole carboxylic acid; SAICAR: 5’-phosphoribosyl-4-(*N*-succino-carboxamide)-5-aminoimidazole; AICAR: 5-aminoimidazole-4-carboxamide ribonucleotide; IMP: inosine monophosphate; XMP: xanthine monophosphate; UMP: uridine monophosphate.

Mutants in genes involved in the synthesis of sulfur-containing amino acids (*cysIJQ, metB*), lysine (lysA) and leucine (*leuABC*) are disadvantaged in chicory (Table 2 and Fig. 3A). These amino acids are known to be present in low concentration in plant tissues. Other amino acids seem to be present in quantity sufficient for growth of *D. dadantii* auxotrophs. Low level of certain amino acids probably induces the stringent response in the bacteria. Reduced growth in the plant of the *relA* mutant, unable to synthesize the alarmone ppGpp, supports this hypothesis. Glucose is the main sugar present in plant tissue, present as a circulating sugar or a cellulose degradation product. Mutants in the PTS glucose transport system genes *ptsI* and *ptsG* have a reduced growth in bacteria (Table 2) showing its importance as a carbon source *in planta*.

**Fig 3.**
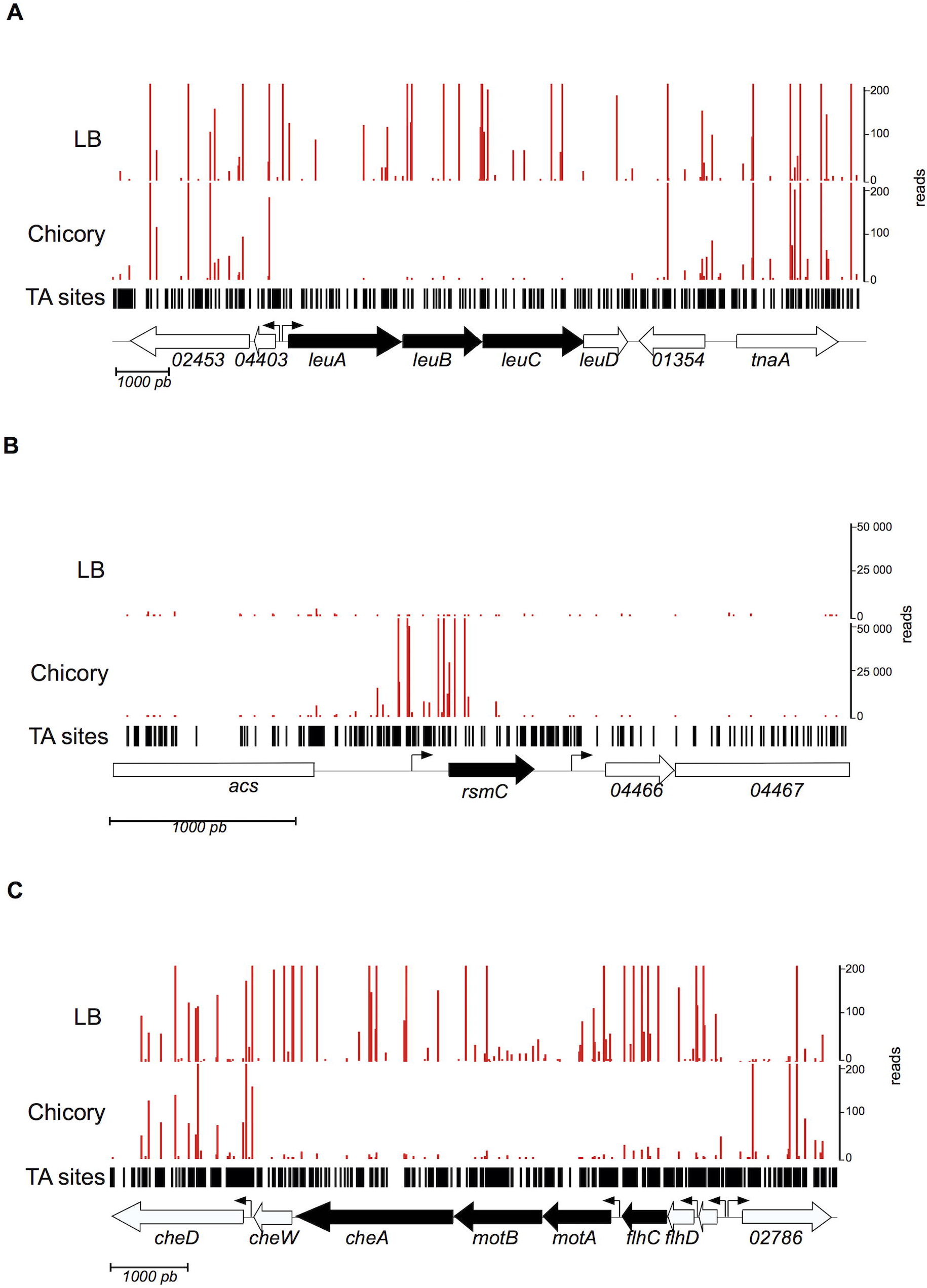
Examples of essential and important genes revealed by Tn-seq. Number of reads at each transposon location in the sample grown in LB or in chicory. Data are averaged from biological replicates and normalized as described in the method section. Three regions of the genome representative of Tn-seq results are shown, with the predicted genes represented at the bottom of each panel. Peaks represent read number at TA sites. Black arrows represent genes that pass the permutation test (q-value ≤ 0.05). Small arrows indicate the presence of promoter (A) Essentiality of leucine biosynthetic genes in chicory. (B) Insertions in the 5’ region of *rsmC* generate growth advantage for the bacteria in chicory. (C) Importance of genes involved in motility for growth in chicory.

Degradation of cell wall pectin by a battery of extracellular enzymes is the main determinant of *Dickeya* pathogenicity. Mutants unable to produce or to secrete these enzymes by the type II secretion system were not disfavored in chicory since these mutants could use for their growth the pectin degradation compounds produced by enzymes secreted by other bacteria. The redundancy of oligogalacturonate specific porins (KdgM and KdgN) and inner membrane transporters (TogT and TogMNABC) allow entry of these compounds into the bacteria even in a mutant in one of these transport systems. However, *kduI* mutants, blocked in the intracellular part of the pectin degradation pathway, have a limited growth *in planta*, confirming the importance of the pectin degradation pathway in the disease progression.

#### (ii) Stress resistance

Plant is an hostile environment for the bacteria that has to cope with antimicrobial peptides, ROS, toxic compounds and acidic pH [35]. We observed that the pump AcrABTolC, that can efflux a wide range of compounds [36], is important for survival in chicory (Fig. S3). Stress can lead to the accumulation of phospholipids in the outer membrane. This accumulation makes the bacteria more sensitive to small toxic molecules [37]. Such a phospholipid accumulation probably occurs when the bacteria infect chicory since *mlaC* and *mlaF* mutants, which are unable to prevent phospholipid accumulation in the outer membrane, have a reduced growth in plant. Production of exopolysaccharides (EPS) was shown to protect the bacteria during the first steps of infection [9]. We observed that *rffG* and *wzx* mutants unable to synthesize EPS have a growth defect in chicory. A set of genes required to repair or degrade altered proteins (*clpA, degQ, trxB*) are also important for survival *in planta*. No gene directly involved in detoxification of ROS was detected in our analysis. However, ROS can create DNA damage. The two helicases involved in DNA repair, UvrD and HelD, give growth advantage in plant. Osmoregulated periplasmic glycans (OPG) are polymers of glucose found in the periplasm of α, β and γ-proteobacteria. Their exact role is unknown but their absence leads to avirulence in certain bacteria such as *D. dadantii* [38]. This absence induces a membrane stress that is sensed and transduced by the Rcs envelope stress response system. This system controls the expression of many genes, including those involved in motility, and those encoding plant cell wall degrading enzymes through the RsmA-RsmB system [39–41]. Thus, mutants defective in OPG synthesis are expected to have a reduced virulence. Indeed, in our experiment, mutants in the two genes involved in OPG synthesis, *opgG* and *opgH* were non competitive in chicory (Table 2).

#### (iii) Iron uptake

*D. dadantii* produces two types of siderophores, achromobactin and chrysobactin, that are required for the development of maceration symptoms in the iron limited environment of plant hosts [42]. Once iron loaded, the siderophores are imported into the bacteria. Import through the outer membrane requires a specific outer membrane channel and the energy transducing complex formed by TonB ExbB and ExbD. While the absence of synthesis of one of the siderophores can be compensated by the presence of siderophore secreted by other bacteria in the growth medium, mutants of the TonB complex are totally unable to acquire iron and thus are unable to grow in the plant. In accordance, *tonB* was essential in chicory while the genes coding for siderophore synthesis or secretion were not. Similarly a mutant devoid of the iron-loaded chrysobactin transport gene (*fct*) is non-competitive.

#### (iv) Regulation

Mutants in several genes controlling virulence factor production have a growth defect in the plant. The master regulator FlhDC acts as a regulator of both flagella and virulence factor synthesis in many bacteria such as *Yersinia ruckeri, Edwardsiella tarda* and *Ralstonia solanacearum* [43–45]. In *D. dadantii* FlhDC has recently been shown to control, in addition to flagellar motility, type III secretion system and virulence factor synthesis through several pathways [46]. We observed that *flhC* gives a growth advantage in chicory. In addition, we uncovered that some genes regulating *flhDC* in other bacteria regulate *D. dadantii* virulence, probably by controlling *flhDC* expression. *rsmC* is a poorly characterized gene in *D. dadantii* but that has been studied in *Pectobacterium carotovorum*. It negatively controls motility and extracellular enzyme production through modulating transcriptional activity of FlhCD [47]. HdfR is a poorly characterized LysR family regulator that controls the *std* fimbrial operon in *S. enterica* and FlhDC expression in *E. coli* [48]. *rsmC* mutants were overrepresented in the chicory (Fig. 3B), indicating a gain of virulence for these mutants. *hdfR* conferred fitness benefits during growth in chicory and could also act in *D. dadantii* as activator of *flhDC* expression.

The GGDEF proteins are c-di-GMP synthase. Their genes are often located next to their cognate EAL diguanylate phosphodiesterase gene. *ecpC (yhjH)* encodes an EAL protein that was shown to activate virulence factor production in *D. dadantii* [49]. *gcpA*, which is located next to *ecpC* encodes a GGDEF protein. Role of *gcpA* in *D. dadantii* virulence has recently been described [50]. We observed that *gcpA* mutants (Dda_03858) were overrepresented in chicory (Table 2). This increased virulence, a phenotype opposite to that described for the *ecpC* mutants, indicates that overproduction of c-di-GMP could reduce *D. dadantii* virulence. Of the eighteen regulators of the LacI family present in *D. dadantii*, four of them were found to be involved in plant infection [51]. One of those, LfcR, which has been found important for infection of chicory, Saintpaulia and *Arabidopsis*, was identified as important for chicory infection in our experiment. LfcR is a repressor of adjacent genes [51]. Surprisingly none of these genes appeared to play a role for chicory infection suggesting that other targets of LfcR probably remain to be discovered.

Finally, it is noteworthy to mention that the *ackA* and *pta* genes are GD *in planta*. These genes constitute the reversible Pta-AckA pathway. The steady-state concentration of acetyl-phosphate (acetyl-P), a signaling molecule in bacteria, depends upon the rate of its formation catalyzed by Pta and of its degradation catalyzed by AckA [52]. The GD phenotype of *D. dadantii ackA* and *pta* mutants during infection suggests that acetyl-P might play a crucial signaling role in the adaptation of *D. dadantii* to the plant tissue.

#### (v) Motility

Motility is an essential virulence factor of *D. dadantii* required to move on the surface of the leaf, enter the wounds and propagate into the plant tissue [53–55]. Accordingly, all the genes required for flagella synthesis, the flagella motor and genes regulating their synthesis (*flhC, flhD, fliA*) (see above) are necessary for fitness during chicory infection (Fig. 3C and 5A). All the genes responsible for the transduction of the chemotaxis signal (*cheA, B, R, W, X, Y* and *Z*) also confer a benefit *in planta* (Table 2). No methyl-accepting chemoreceptor gene mutant was found. Like other environmental bacteria, *D. dadantii* encodes many such proteins (47). They probably present some redundancy in the recognized signal which prevented their detection by our screen.

### *D. dadantii* flagellin is modified by glycosylation

A group of six genes located between *fliA* and *fliC* retained our interest since insertions in these genes led to a growth defect in chicory (Fig. 4A). This effect does not result from insertions in the first gene of the group since they are not expressed in operon [56]. Dda3937_03424 encodes an O-linked N-acetylglucosamine transferase and Dda3937_03419 encodes a protein with a nucleotide diphospho sugar transferase predicted activity. The other ones could be involved in the modification of sugars (predicted function of Dda3937_03423: nucleotide sugar transaminase, Dda3937_03422: carbamoyl phosphate synthase, Dda3937_03421: oxidoreductase; Dda3937_03420: methyltransferase). Their location let suppose that this group of genes could be involved in flagellin glycosylation. Analysis by SDS-PAGE of FliC produced by the wild type, and the Dda3937_03424 and Dda3937_03419 mutants, showed that in the two latter strains the molecular weight of the protein diminished (Fig. 4B). The molecular weight determined by mass spectroscopy was 28,890 Da for FliC_A42_77, 31,034 Da for FliC_A3422_ and 32170 Da for the WT FliC. Thus, in the wild type strain FliC is modified by the products of the genes Dda_03424 to Dda_03419, probably by multiple glycosylation with a disaccharide. Absence of modification did not modify *D. dadantii* motility (data not shown). The flagellin of the plant pathogens *Pseudomonas syringae* pv *tabaci* and *Burkholderia cenocepacia* are also glycosylated and absence of this modification lowered the ability of these bacteria to cause disease on tobacco and *Arabidopsis*, respectively [57, 58]. Accordingly, in *D. dadantii*, FliC modification appears important for multiplication of the bacteria in plant.

**Fig 4.**
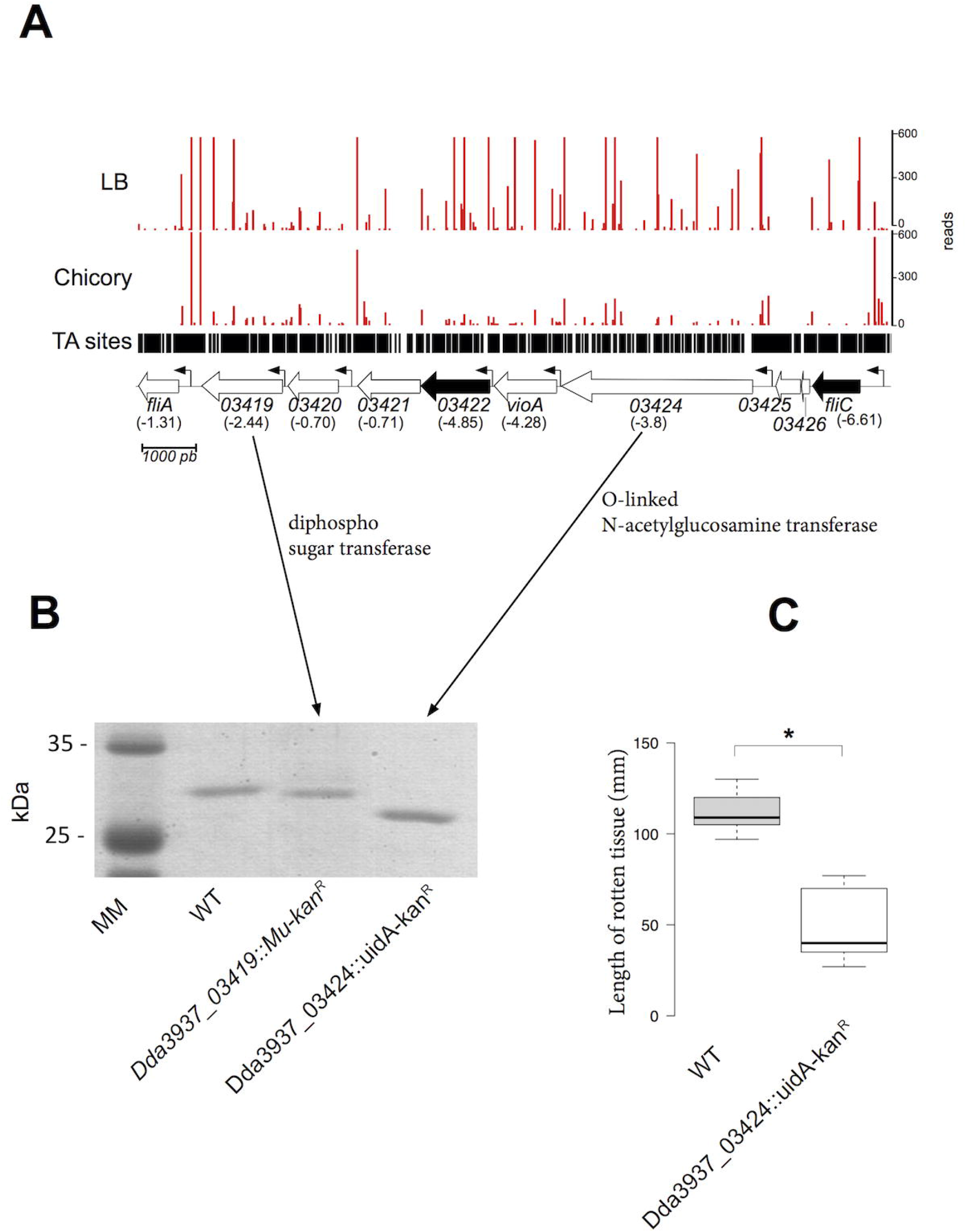
Modification of FliC revealed by Tn-seq analysis and SDS-PAGE. (A) Importance of 6 genes located between *fliA* and *fliC* for growth in chicory. Log_2_FC are indicated in bracket. Dda3937_03425 and Dda3937_03426 are duplicated transposase genes that have been removed from the analysis. Black arrow: GD in chicory (q-value ≤ 0.05); white arrow: genes that do not pass the permutation test (q-value > 0.05). Small arrows indicate the presence of promoter. (B) Analysis by SDS-PAGE of FliC produced by the wild type (lane 1), the A3422 (lane 2) and A4277 (lane 3) strains. (C) Maceration of celery leaves by the Wild Type and A4277 (glycosylation) mutant. Length of rotten tissue was measured 48 h post infection. Boxplot were generated by BoxPlotR from 9 data points. The calculated median value is 109 for the WT strain, 40 for the A4277 strain. Center lines show the medians; box limits indicate the 25th and 75th percentiles as determined by R software; whiskers extend 1.5 times the interquartile range from the 25th and 75th percentiles.

**Fig 5.**
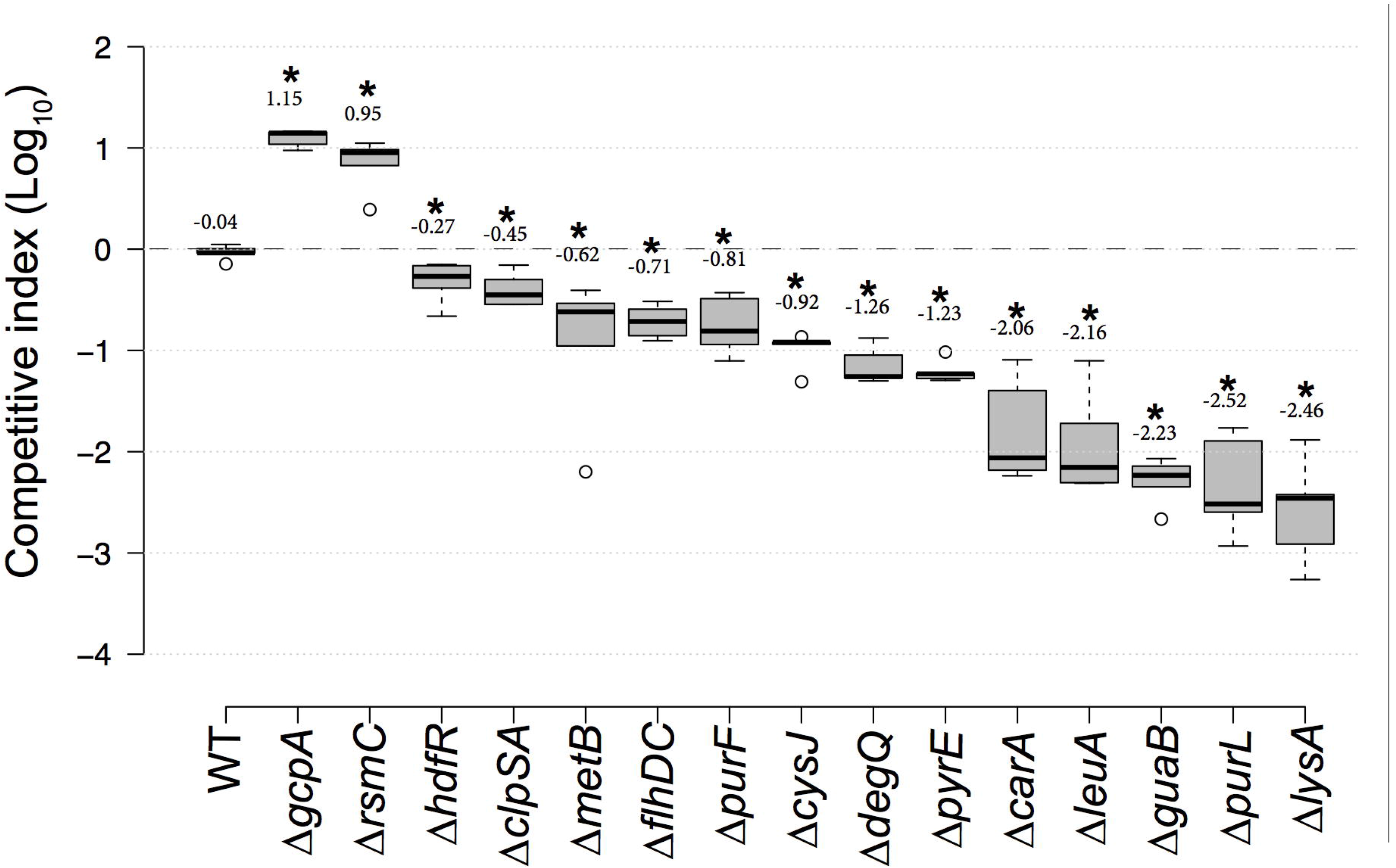
Competitive Index (CI) of several mutant strains. CI values were determined in chicory leaves as described in Methods. Each value is the mean of 5 experiments. Center lines show the medians; box limits indicate the 25th and 75th percentiles as determined by R software; whiskers extend 1.5 times the interquartile range from the 25th and 75th percentiles, outliers are represented by dots. n = 5 sample points. Numbers above the boxes indicate the average competitive index in Log_10_. * indicates a significant difference relative to the WT (p<0.05, Welch’s t-test).

### Additional genes could be involved in virulence

Several genes have a log_2_FC >4 or <−4 but do not satisfy the statistical permutation test adjusted by the false discovery rate method (q-value) (table S6). However, most of them belong to the categories described above and could be required for growth *in planta*. Among those with a log_2_FC< −4 can be found genes involved in amino acid and nucleic acid synthesis (*cysH, ilvC, pyrF, pyrD, purC, thrC, metA, cysK, lysC*), flagella and motility *flgJ, fliO, flgC, fliS, flgG, flgA, flgL, cheW, fliN, fliP, fliK, fliG, fliL*), pectin and glucose metabolism (*kduD, pgi*), EPS synthesis (*gmd*), flagella glycosylation (vioA) and regulation (zur, *ecpC* and the general RNA chaperone *hfq*).

Among the genes with a log2FC > 4, three regulators can be noticed: *pecS*, *pecT* and *Dda3937_00840. pecS* and *pecT* are known regulators of *D. dadantii* controlling the expression of many factors involved in virulence [5, 6]. Thus, their mutation could confer an increased fitness of the bacteria in chicory. *D. dadantii* possesses a functional *expI-expR* quorum sensing system which does not seem to control plant virulence factor production [59]. However, several LuxR family regulator genes which are not associated with a *luxI* gene are present in the genome of the bacteria. Mutants of one of them (*Dda3937_00840*) are overrepresented in the chicory. Its product is probably a repressor of genes conferring an increased fitness *in planta*.

### Validation of the Tn-seq results

To validate the Tn-seq results, we performed coinoculation experiments in chicory leaves with the wild type strain and various mutants in GA genes (*gcpA* and *rsmC*) or GD genes (*hdfR, clpSA, metB, flhDC, purF, cysJ, degQ, pyrE, carA, leuA, guaB, purL, lysA*) in a 1/1 ratio. We calculated a competitive index (CI) by counting the number of each type of bacteria in the rotten tissue after 24 h. We confirmed the ability of the *ΔrsmC* and *ΔgcpA* to overgrow the wild type strain. At the opposite, the wild type strain overgrew the in frame deletion mutants that were tested. The lowest competitive index were observed with mutants in biosynthetic pathways such as *ΔleuA, ΔguaB, ΔpurL, ΔlysA*.

Amino acid auxotroph mutants (Cys^−^, Leu^−^, Met^−^ and Lys^−^) tested by coinoculation experiments could be phenotypically complemented *in planta*. Addition of both the non-synthetized amino acid and the auxotroph mutant within the wound totally or almost completely suppressed the growth defect of the auxotroph mutant *in planta* (Fig. S4). This confirmed the low availability of certain amino acids in chicory. This result confirmed that Tn-seq is a reliable technique to identify genes involved in plant colonization and virulence.

## Conclusion

This Tn-seq experiment highlights new factors required for *D. dadantii* successful rotting of chicory. Many genes known to be important for pathogenesis were not found in this screen because their products are secreted and can be shared with other strains in the community. This includes all the proteins secreted by the type II secretion system and small molecules such as siderophores and butanediol. Other categories of genes were not found: for example, no genes involved in response to acidic or oxidative stresses were identified. Chicory has been described as an inadequate model to study the response of *D. dadantii* to oxidative stress [60]. Similarly, the type III *hrp* genes were not identified in our study. The Hrp system is not necessary for *D. dadantii* virulence and in our experimental conditions (high inoculum on isolated chicory leaves) the necrotrophic capacities of *D. dadantii* (production of plant cell wall degrading enzymes) is probably sufficient to provoke the disease. Our results also uncover some unknown aspects of the infection process. Struggle for iron availability between plant and bacterial pathogens has been well described. However, a competition for amino acids and nucleic acid seems also to take place in the plant. The amount of nucleic acids and of the cysteine, leucine, methionine, threonine and isoleucine amino acids is too low in chicory to allow an efficient multiplication of bacteria defective in their biosynthesis. Some enzymatic steps involved in their synthesis are specific to bacteria and fungi. Thus, they could be good targets for the development of specific inhibitors [61] to fight *D. dadantii*. Regulation of *D. dadantii* virulence has been extensively studied [2, 21]. However, new regulatory genes controlling virulence were also detected in this study. An orphan LuxR family regulator seems to play an important role in virulence. New members of the FlhDC regulation pathway were also detected. A few genes of unknown function remain to be studied.

*D. dadantii* can infect dozens of plants. Besides chicory, *D. dadantii* virulence tests are usually performed on potato plant, tuber or slices, *Arabidopsis thaliana*, saintpaulia or celery. Metabolic status or reaction defenses of these model plants are all different and bacterial genes required for a successful infection will probably differ in each model. Testing several of them will allow to determine the full virulence repertoire of the bacteria.

While Tn-seq has been used to study genes required for the infection of animals, there has been no genome-wide study of factors necessary for a necrotrophic plant pathogen to develop and provoke disease on a plant. Besides the genes of known function described in the Result section, this study allowed the identification of several genes of unknown function required for chicory rotting. Repetition of this experiment with other strains or on other plants will tell if these genes encode strain or host specific virulence factors.

## Methods

### Bacterial strains and growth conditions

Bacterial strains, phages, plasmids and oligonucleotides used in this study are described in Table S1 to Table S3. *D. dadantii* and *E. coli* cells were grown at 30 and 37°C respectively in LB medium or M63 minimal medium supplemented with a carbon source (2 g/L). When required antibiotics were added at the following concentration: ampicillin, 100 μg/L, kanamycin and chloramphenicol, 25 μg/L. Media were solidified with 1.5 g/L agar. Transduction with phage PhiEC2 was performed according to [62]. The motility of each mutant was compared with that of the wild-type strain on semisolid (0.4%) LB agar plates as previously described [63].

### Construction of the transposon library

Five mL of an overnight culture of *D. dadantii* strain A350 and of *E. coli* MFDpir/pSamEC were mixed and centrifuged 2 min at 6000 g. The bacteria were resuspended in 1 mL of M63 medium and spread onto a 0.45 μm cellulose acetate filter placed on a M63 medium agar plate. After 8h, bacteria were resuspended in 1 mL M63 medium. An aliquot was diluted and spread onto LB agar + kanamycin plates to estimate the efficiency of mutagenesis. The other part was inoculated in 100 mL of LB medium + kanamycin and grown for 24 h at 30°C. To confirm that the bacteria that grew were *D. dadantii* strains with a transposon but without plasmid pSamEC, we checked that all the grown bacteria were kan^R^, amp^S^ and diaminopimelate (DAP) prototrophs (MFDpir is DAP^−^). The bacteria were frozen in 40% glycerol at −80°C and represent a library of about 300 000 mutants.

### DNA preparation for high-throughput sequencing

An aliquot of the mutant library was grown overnight in LB medium + kanamycin. To identify essential genes in LB, the culture was diluted 100-fold in LB and grown for 6 h. To infect chicory, the overnight culture was centrifuged and resuspended at OD_600_ = 1 in M63 medium. Chicories cut in half were inoculated with 10 μL of this bacterial suspension and incubated at 30°C with maximum moist. After 60 h, the rotten tissue was collected and filtered through a cheesecloth. The bacteria were collected by centrifugation and washed twice in M63 medium. DNA was extracted from 1.5 mL aliquots of bacterial suspension adjusted to OD_600_1.5 with the Promega Wizard Genomic DNA purification kit. Next steps of the DNA preparation methods were adapted from [26]. All DNA gel-extraction were performed onto a blue-light transilluminator of DNA stained with gel-green (Biotium) to avoid DNA mutation and double-stranded breaks. 50 μg of DNA samples were digested with 50 U MmeI in a total volume of 1.2 mL for one hour at 37°C according to manufacturer’s instructions, then heat-inactivated for 20 minutes at 80°C, purified (QIAquick, PCR purification kit Qiagen) and concentrated using a vacuum concentrator to a final volume of 25 μL. Digested DNA samples were run on a 1% agarose gel, the 1.0–1.5 kb band containing the transposon and adjacent DNA was cut out and DNA was extracted from the gel according to manufacturer’s instructions (Qiaquik Gel Extraction Kit, Qiagen). This allowed recovery of all the fragments containing genomic DNA adjacent to transposons (1201 bp of transposable element with 32-34 bp of genomic DNA). A pair of single-stranded complementary oligonucleotides containing an unique 5-nt barcode sequence (LIB_AdaptT and LIB_AdaptB) was mixed and heated to 100°C, then slowly cooled down in a water bath to obtain double-stranded adaptors with two-nucleotide overhangs. 1 μg DNA of each sample was ligated to the barcoded adaptors (0.44 mM) with 2000 U T4 DNA ligase in a final volume of 50 μL at 16°C overnight. Five identical PCR reactions from the ligation product were performed to amplify the transposon adjacent DNA. One reaction contained 100 ng of DNA, 1 unit of Q5 DNA polymerase (Biolabs), 1X Q5 Buffer, 0.2 mM dNTPs, 0.4 μM of the forward primer (LIB_PCR_5, which anneals to the P7 Illumina sequence of the transposon) and the reverse primer (LIB_PCR_3, which anneals to the P5 adaptor). Only 18 cycles were performed to keep a proportional amplification of the DNA. Samples were concentrated using a vacuum concentrator to a final volume of 25 μL. Amplified DNA was run on a 1.8% agarose gel and the 125 bp band was cut-out and gel extracted (QIAquick, PCR purification kit Qiagen). DNA was finally dialysed (MF-Millipore™ Membrane Filters) for 4 hours. Quality control of the Tn-seq DNA libraries (size of the fragments and concentration) and High-throughput sequencing on HiSeq 2500 (Illumina) was performed by MGX (CNRS sequencing service, Montpellier). 6 DNA libraries were multiplexed on one flow-cell. After demultiplexing, the total number of reads was comprised between 18 and 31 millions (Table 1).

### Bioinformatics analysis

Raw reads from the fastQ files were first filtered using cutadapt v1.11 [64] and only reads containing the *mariner* inverted left repeat (ACAGGTTGGATGATAAGTCCCCGGTCTT) were trimmed and considered *bona fide* transposon-disrupted genes. Trimmed reads were then analyzed using a modified version of the TPP script available from the TRANSIT software v2.0.2 [32]. The mapping step was modified to select only reads mapping uniquely and without mismatch in the *D. dadantii* 3937 genome (Genbank CP002038.1). Then, the counting step was modified to accurately count the reads mapping to each TA site in the reference genome according to the Tn-seq protocol used in this study. Read counts per insertion were normalized using the LOESS method as described in [65]. We next used the TRANSIT software (version 2.0) to compare the Tn-seq datasets.

### Strain construction

To construct the A4277 strain, gene Dda3937_03424 was amplified with the oligonucleotides 19732+ and 19732−. The resulting fragment was inserted into the pGEM-T plasmid (Promega). A uidA-kan^R^ cassette [66] was inserted into the unique AgeI site of the fragment. The construct was recombined into the *D. dadantii* chromosome according to [67]. Recombination was checked by PCR. To construct the in-frame deletion mutants, the counter-selection method using the *sacB* gene was used [68]. The suicide pRE112 plasmid containing 500 bp of upstream and downstream DNA of the gene to delete was transferred by conjugation from the *E. coli MFDpir* strain into *D. dadantii* 3937. Selection of the first event of recombination was performed on LB agar supplemented with chloramphenicol at 30 μg/L. Transconjugants were then spread on LB agar without NaCl supplemented with 5 % sucrose to allow the second event of recombination. In-frame deletions were then checked by auxotrophy analysis and/or by PCR (Dreamtaq polymerase, Thermofisher). In order to discriminate mutants from the wild strain during coinoculation experiments, a Gm^R^ derivative of the WT strain was constructed by insertion of the mini-Tn7-Gm into the *attTn7* site (close to the *glmS* gene) [69]. A 3937 Gm^R^ strain was made by coelectroporation of pTn7-M [69] and pTnS3 [70] plasmids into *D. dadantii* 3937 strain. The mini-Tn7-Gm delivered by the pTn7-M vector (suicide plasmid in *D. dadantii*) is inserted into the *attTn7* site (close to the *glmS* gene) of recipient strain thanks to pTnS3 plasmid encoding the Tn7 site-specific transposition pathway. The Gm^R^ strain obtained was then checked by PCR using attTn7-Dickeya3937-verif and 3-Tn7L primers (Table S3).

### Protein techniques

Flagella were prepared from overnight LB grown cells. Bacteria were pelleted, resuspended in 1/10 volume of water and passed 20 fold through a needle on a syringe. Cells and cells debris were removed by centrifugation 5 min at 20 000 × g [63]. Proteins were analyzed by SDS-polyacrylamide gel electrophoresis (SDS-PAGE).

### Celery inoculation experiments

Wild Type and A4277 (glycosylation) mutant were grown overnight in M63 + glycerol medium. Bacteria were washed in M63 medium and the OD_600_ was adjusted to 1.0. Bacteria were diluted 10-fold in the same medium. 10 μL of the bacterial suspension were inoculated into leaves in a hole made with a pipet tip. The wound was covered with mineral oil and the leaves were incubated at 30°C at high humidity for 2 days (celery). Length or rotten tissue was measured.

### Coinoculation experiments

To determine the competitive index of the mutants, the wild type strain and the mutant to test were grown overnight in M63 + glycerol medium. Bacteria were washed in M63 medium and the OD_600_ was adjusted to 1.0. Bacteria were mixed to a 1:1 ratio and diluted 10-fold. For complementation experiments *in planta*, the dilution was performed in M63 medium with 1mM of required amino acid. 10 μL of the mixture were inoculated into chicory leaves. The wound was covered with mineral oil and the leaves were incubated at 30 °C at high humidity. After 24 h the rotten tissue was collected, homogenized, diluted in M63 and spread onto LB and LB + antibiotic plates. After 48 h at 30°C, colonies were counted. The competitive index is the ratio (number of mutant bacteria/number of WT bacteria) in the rotten tissue / (number of mutant bacteria/number of WT bacteria) in the inoculum. For the genes whose absence confers a growth advantage in chicory according to the Tn-seq experiment, in frame deletions were realized in a WT strain. The other mutants were realized in the 3937 Gm^R^ strain.

### Nucleotide sequence accession numbers

The transposon sequence reads we obtained have been submitted to the ENA database under accession number PRJEB20574.

## Acknowledgments

We thank Geraldine Effantin, Veronique Utzinger, Matthias Schulz, Andrea Flipo, Leana Corneloup and Barbara Gbaguidi for technical assistance, the members of the MTSB team and Xavier Charpentier for discussion, Nicole Cotte-Pattat and Sarah Bigot for critical read of the manuscript and James Paslesvert for encouragements.

## Supporting information legends

**Fig S1. Biological reproducibility of the Tn-seq results**. Pairs of Tn-seq assay results are compared, with the total number of reads per gene plotted. Analysis of DNA samples corresponding to two independent cultures of the mutant pool grown (A) in LB medium (correlation coefficient R = 0.72) and (B) in chicory (correlation coefficient R = 0.98). Values represent average numbers of reads per gene from the pairs of biological replicates.s

**Fig S2. Volcano plot of resampling results comparing replicates grown in chicory versus in LB**. Significant hits have q < 0.05 or −log_10_ q > 1.3. Growth defect (GD) or growth advantage (GA) genes are indicated by a red frame.

**Fig S3. *acrAB* are essential in chicory**. Number of Tn-seq reads at each insertion site in the *acrA acrB* region in samples grown in LB or in chicory. Data are averaged from biological replicates and normalized as described in Methods. *dnaX* which encodes both the tau and gamma subunits of DNA polymerase is represented by a grey arrow. *dnaX* is essential gene in LB. *acrAB* represented by grey arrows are GD in chicoy (q-value ≤ 0.05).

**Fig S4. Complementation of auxotroph mutants *in planta***. Each leaf was inoculated with 10^6^ bacteria. Length of rotten tissue was observed after 24h. Bacteria were injected into the wounded leaf with or without amino acid. Center lines show the medians; box limits indicate the 25th and 75th percentiles as determined by R software; whiskers extend 1.5 times the interquartile range from the 25th and 75th percentiles, outliers are represented by dots. n = 5 sample points. Numbers above the boxes indicate the average competitive index in Log_10_. * indicates a significant difference relative to the WT (p<0.05, Welch’s t-test). ** Indicates an absence of significant difference relative to the WT (p>0.05, Welch’s t-test).

**Table S1: bacterial strains used in this study**

**Table S2: plasmids used in this study**

**Table S3: oligonucleotides used in this study**

**Table S4: number of genes implicated in KEGG pathway**

**Table S5: raw data of the HMM and resampling analysis by transit**

**Table S6: List of genes with log_2_FC <−2 or >2 but with q-value >0.05**

